# Production of santalenes and bergamotene in *Nicotiana tabacum* plants

**DOI:** 10.1101/398776

**Authors:** Jun-Lin Yin, Woon-Seng Wong

## Abstract

Terpenes play an important role in plant–insect relationships, and these relationships can potentially be modified by altering the profile of terpenes emitted from plants using metabolic engineering methods. Transgenic plants generated by employing such methods offer the prospect of low-cost sustainable pest management; in this regard, we used chloroplast targeting and cytosolic mevalonic acid pathway enhancement in this study to investigate the interaction of santalenes and bergamotene with insects. The santalene- and bergamotene-emitting transgenic tobacco plants thus generated were utilized to study host preference in the green peach aphid *Myzus persicae* (Sulzer). The results showed that co-expression of either 3-hydroxy-3-methylglutaryl-CoA reductase (HMGR) or truncated HMGR with santalene synthase led to the production of higher amounts of santalenes and bergamotene in transgenic tobacco plants, and that these santalene- and bergamotene-emitting plants were attractive to green peach aphids. We accordingly propose that such transgenic plants may have potential application in pest management as a trap crop to prevent green peach aphid infestation of wild-type tobacco plants.

## Introduction

Volatile terpenoids play important roles in plant defense, as they normally act as deterrents or repellents against herbivores, and can also function as attractants to attract predators [1-6]. Although the application of insecticides is a common and effective method of pest management, these chemicals can have undesirable side effects on the environment, promote rapid evolution of resistance in pests, and can reduce predator populations [7]. Previous studies have demonstrated that transgenic plants in which emitted volatiles had been modified by metabolic engineering can be used to deter feeding by hornworms [8], decrease aphid colonization [9-11], or attract mite predators of infesting arthropods [12]. Modification of plant volatile terpenoids through metabolic engineering thus has considerable potential to lower the cost of pest management and facilitate more sustainable agricultural practices by reducing reliance on chemical pesticide usage [13].

Volatile terpenes, including monoterpenes (C_10_), sesquiterpenes (C_15_), and diterpenes (C_20_), play an important role in a wide range of plants. Plants interact with insects, surrounding plants, and pathogens by releasing volatile molecules. Monoterpenes (C_10_) and diterpenes (C_20_) are mainly biosynthesized via the plastidial methylerythritol 4-phosphate (MEP) pathway, while sesquiterpenes (C_15_) are mainly biosynthesized from isopentenyl diphosphate (IPP) and dimethylallyl diphosphate (DMAPP) via the cytosolic mevalonic-acid (MVA) pathway.[14]. In recent years, *Nicotiana tabacum* has served as a platform for the production of several sesquiterpenes, as this plant can be sustainably cultivated at low cost [15]. Several specific metabolic engineering methods have been developed to enhance the production of sesquiterpenes in tobacco. For instance, the reaction catalyzed by 3-hydroxy-3-methylglutaryl-CoA reductase (HMGR) is believed to be an important rate-limiting step in the tobacco MVA pathway; tobacco plants over-expressing *HMGR* have been shown to accumulate three to six times more sterol than wild-type plants [16]. In addition to enhancing *HMGR* expression, transgenic tobacco plants in which both sesquiterpene synthase and farnesyl diphosphate synthase (FPS) had been targeted into chloroplasts produced 1000-fold more patchoulol or amorpha-4,11-diene compared with transgenic lines in which the responsible sesquiterpene synthase gene was expressed only in the cytosol [8,17]. Co-expression of the mitochondrial *amorpha-4,11-diene synthase* (*ADS*) gene and a truncated *HMGR* (*tHMGR*) gene has been found to increase the production of amorpha-4,11-diene in transgenic tobacco [18,19], whereas transgenic tobacco co-expressing *tHMGR* and *FPS* with a *valencene synthase* gene showed increased production of (+)-valencene [20]. Furthermore, squalene biosynthesis was influenced by engineering genes of FPS and squalene synthase (SQS) via the *N. tabacum* chloroplast genomes [21]. Therefore, co-expression of HMGR, subcellular localization of sesquiterpene synthase, and chloroplast genome transformation are considered to be three important factors that determine the production of sesquiterpenes in transgenic tobacco plants.

Sandalwood is an essential oil that is extracted from the heartwood of sandalwood plants (*Santalum album)* and is used as a popular fragrance and insect repellent [22]. However, given that sandalwood is a rare resource, the utilization of sandalwood oil is somewhat limited. Santalols and bergamotol are the major constituents of this essential oil, and are generated from santalene/bergamotene by santalene/bergamotene oxidases. In *S. album* plants, santalenes and bergamotene can be generated from farnesyl diphosphate (FPP) in the MVA pathway via the activity of santalene synthase (SaSSy) [23,24]. Green peach aphid (*Myzus persicae* (Sulzer)), an important pest affecting tobacco production, has a broad spectrum of hosts worldwide [25,26]. To date, however, metabolic engineering of santalenes and bergamotene in tobacco plants, and their interaction with green peach aphid have not been intensively studied. In the present study, we employed chloroplast targeting methodology and co-expression of FPS and HMGR or tHMGR to enhance the production of santalenes and bergamotene in *Nicotiana benthamiana* and *N. tabacum* plants. The results showed that co-expression of either *HMGR* or *tHMGR* with *SaSSy* enhanced the production of santalenes and bergamotene in both these tobacco species. In addition, to optimize biosynthesis of santalenes and bergamotene in tobacco plants, we investigated insect interaction of these sesquiterpenes in green peach aphid choice experiments. In both short-term and long-term choice experiments, transgenic tobacco plants emitting santalenes and bergamotene showed increased attraction to the green peach aphid *M. persicae* (Sulzer), indicating that these transgenic tobacco plants have considerable potential as a trap crop to protect wild-type tobacco from *M. persicae* feeding damage.

## Materials and methods

### Plant materials and growth conditions

The seeds of tobacco plants (*N. benthamiana* and *N. tabacum* SR1) provided by Prof. Nam-Hai Chua (The Rockefeller University), were grown in a glasshouse under natural light conditions. Final binary vectors were introduced into *Agrobacterium tumefaciens* AGL1 by electro-transformation, and the transformed *A. tumefaciens* were then infiltrated into the leaves of tobacco plants using a standard protocol [16].

### Vector construction

All the vectors used in this study are shown in S1 Fig. Proof HF MasterMix (Bio-Rad, USA) and Rapid DNA ligation kit (ROCHE, Swiss) were used to perform PCR and ligation, respectively.

Sequences of the RbcS secretion signal (*RTP*) (GenBank: NM_202369.2) and a linalool synthase transit peptide (*LTP*) (GenBank: JN408286.1) were amplified from *Arabidopsis thaliana* cDNA and *Solanum lycopersicum* cDNA, respectively. These sequences were then ligated into pGEM-T Easy vectors (Promega) and named pGEM-RTP and pGEM-LTP, respectively. *SaSSy* and *FPS* (GenBank: HQ343276.1 and U80605.1) were amplified and ligated into the pGEM-RTP and pGEM-LTP vectors to form pGEM-RTPFPS, pGEM-RTPSaSSy, pGEM-LTPFPS, and pGEM-LTPSaSSy constructs. The *RTPSaSSy, RTPFPS, LTPSaSSy*, and *LTPFPS* fragments were subsequently inserted into pCAMBIA1300-GFP vectors, yielding 1300-35S-RTPSaSSy-GFP, 1300-35S-RTPFPS-GFP, 1300-35S-LTPSaSSy-GFP, and 1300-35S-LTPFPS-GFP constructs, respectively. Furthermore, *SaSSy* or *FPS* was amplified and ligated into pCAMBIA1300-GFP vectors to generate 1300-35S-SaSSy-GFP and 1300-35S-FPS-GFP, respectively. The 35S promoter of the 1300-35S-RTPSaSSy-GFP and 1300-35S-SaSSy-GFP vectors was subsequently replaced with the *A. thaliana RbcS* promoter (GenBank: X13611.1) to yield the vectors 1300-RbcSP-RTPSaSSy-GFP and 1300-RbcSP-SaSSy-GFP, respectively.

*tHMGR* and *HMGR* (GenBank: AY488113.1) were amplified from *A. thaliana* cDNA and cloned into pSAT1A vectors. The *OCS* promoter of the pSAT1A vectors was replaced with the *ubiquitin 10* promoter [27]. UBQP-tHMGR-OCST and UBQP-HMGR-OCST fragments were ligated into the 1300-RbcSP-SaSSy-GFP vector, to generate 1300-tHMGR-SaSSy and 1300-HMGR-SaSSy vectors, respectively.

*RTPFPS, FPS,* and *SaCYPF39v1* (GenBank: KC533716.1, synthesized by Genscript) were cloned into pBA002 vectors to generate pBA-RTPFPS, pBA-FPS, and pBA-SaCYPF39v, respectively. *tHMGR* and *HMGR* were also cloned into pB7WG2D vectors to generate pB7WG2D-tHMGR and pB7WG2D-HMGR, respectively, via Gateway cloning (Invitrogen, USA). All primers, genes, and vectors used in these procedures are shown in S1 and S1 Tables, and S1 Fig.

### Infiltration of *nicotiana benthamlana* leaves

*A. tumefaciens* cells were cultured in LB medium containing kanamycin (25 mg/L), spectinomycin (50 mg/L), or refampicillin (50 mg/L) and incubated at 28 °C at 250 rpm for 2 d. Thereafter, the cells were centrifuged for 10 min at 4500 rpm and 4 °C, resuspended with 10 mM MES buffer (containing 10 mM MgCl_2_ and 100 μM acetosyringone) to a final OD_600_ of 1.0, and then incubated at room temperature for 2 h. An *A. tumefaciens* strain encoding the TBSV P19 protein was added to suppress gene silencing. Using a 1-mL syringe, *A. tumefaciens* cells were infiltrated into 3-week-old *N. benthamiana* plants grown in a glasshouse under natural light conditions. Three days after infiltration, the terpene products of the treated plants were analyzed by gas chromatography–mass spectrometry (GC–MS).

### Protein location

The *A. tumefaciens* strain AGL1, carrying 1300-RTPSaSSy-GFP, 1300-RTPFPS-GFP, 1300-LTPSaSSy-GFP, or 1300-LTPFPS-GFP plasmids, were injected into *N*. *benthamiana* leaves. The *A. tumefaciens* strain encoding the TBSV P19 protein was added to suppress gene silencing. After 2–4 days, protein localization was observed by confocal laser scanning microscopy.

### Analyses of extracts and headspace volatiles

Two hundred milligrams of infiltrated leaves was ground in liquid nitrogen and extracted with 0.5 mL ethyl acetate at room temperature. One microliter of camphor–methanol solution (10 mg/mL), which was used as an internal standard, was added to the extracts. The extracts were prepared by shaking for 1 h at room temperature. Thereafter, the extracts were centrifuged for 5 min at 12000 rpm, and the upper organic layer was transferred to a new tube and dehydrated using anhydrous sodium sulfate (Na_2_SO_4_). GC– MS was used to analyze the extracts. Volatiles were collected from 10 g of detached transgenic tobacco leaves using a Push–pull headspace collection system [28] under standard growth chamber conditions (25 °C, 16-h light:8-h dark photoperiod). Two microliters of camphor–methanol solution (10 mg/mL), which was used as an internal standard, was added to the leaf surfaces. A Porapak Q trap (80–100 mesh size; Sigma, USA) was used to trap volatiles during a 24-h period. Volatiles were eluted two times with 200 μL of dichloromethane and analyzed using an Agilent GC 7890A system incorporating a 5975C inert mass-selective detector and an HP-5 MS column (30 m × 0.25 mm; 0.25 μm film thickness). Samples (2 μL) were injected at an initial oven temperature of 50 °C, which was held for 1 min and then subsequently increased to 300 °C at a rate of 8 °C/min and then held for 5 min. Compounds were identified by comparison of their mass spectra with those in the NIST MS 2014 library.

### RNA isolation and quantitative real-time PCR

Total RNA was isolated from transgenic tobacco using an RNeasy Plant Mini Kit (Qiagen, German), following the manufacturer’s instructions, and cDNA was synthesized using iScript RT SuperMix (Bio-Rad, USA). Gene-specific primers (shown in Table S3) were designed using Primer3Plus. Quantitative PCR was performed using Power SYBR^®^ Green PCR Master Mix (Life Technology, USA) in an ABI 7900 RT-PCR system (Applied Biosystems, USA). The transcript levels of all genes were analyzed relative to a reference gene, elongation factor-1 alpha (*EF1a*; GenBank: D63396.1), using the Δ Δ Ct method [29]. For each experiment, three replicates were performed with at least three independent biological samples.

### Aphids

Green peach aphids (*Myzus persicae* [Sulzer]) were isolated from Singapore and reared in a 52 × 52 × 50 cm cage containing eight *A. thaliana* plants and maintained in a growth chamber (25 °C, 16-h light:8-h dark photoperiod).

### Short- and long-term Insect choice experiments using transgenic tobacco

A short-term choice experiment using green peach aphids was modified from that described previously [30]. Two plants of similar size and leaf number (one wild-type [WT] and one transgenic tobacco plant) were placed in a cage (30 × 30 × 30 cm), into which 30 adult green peach aphids were released. Twenty-four hours after release, the number of aphids on each plant was recorded.

To investigate the long-term choice preferences of these aphids, two plants of similar size and leaf number (one WT and one transgenic tobacco plant) were grown in a cage (30 × 30 × 30 cm) (25 °C, 16-h light:8-h dark photoperiod), into which 30 adult green peach aphids were released. Four weeks after release, the number of aphids on each plant was recorded.

### Aphid two-choice experiments using hexane solutions of santalenes and bergamotene

Two leaves from 3-week-old WT *N. tabacum* plants were placed on the surface of agar medium in plates, one leaf was treated with 20 µL of hexane solutions of santalenes and bergamotene hexane (1 gL^-1^), and the other leaf was treated with 20 µL of hexane as a control. After evaporation of the solvent, 10 green peach aphids were released in each plate, and 15 min after release, the number of aphids on or above the leaf was recorded.

### Statistical analysis

Values are indicated as ‘mean ± SD’ for three to six biological replicates. Statistical comparison was analyzed using a two-tailed Student’s *t*-test or unequal variance *t*′-test, and indicated by asterisks. * indicates P < 0.05; ** indicates P < 0.01.

## Results

### Targeting of FPS and SaSSy into chloroplasts using chloroplast transit peptides

Most chloroplast proteins are encoded by the nuclear genome and synthesized as precursors containing N-terminal targeting transit peptides. A chloroplast target peptide is a short peptide chain that directly transports a protein to a chloroplast, after which most of the transit peptide is cleaved from the protein [31]. Targeting efficiency is affected by the structure and sequence motifs of the transit peptide [32]. Sesquiterpenes are naturally synthesized from IPP and DMAPP via the cytosolic MVA pathway [14]. In order to synthesize sesquiterpenes in chloroplasts, we used a *S. lycopersicum* linalool synthase chloroplast transit peptide (LTP) and an *A. thaliana* ribulose bisphosphate carboxylase small subunit chloroplast transit peptide (RTP) to target both *A. thaliana* FPS and SaSSy proteins into chloroplasts.

*FPS, SaSSy, LTP,* and *RTP* sequences were inserted into pCambia1300-GFP vectors to generate 1300-35SP-LTP-FPS-GFP, 1300-35SP-LTP-SaSSy-GFP, 1300-35SP-RTP-FPS-GFP, and 1300-35SP-RTP-SaSSy-GFP vectors, respectively (S1 Fig.). Three days after *N. benthamiana* agro-infiltration, the subcellular locations of FPS and SaSSy proteins were examined using confocal microscopy (Fig. 1). The presence of a green fluorescent protein signal (GFP) showed that native FPS and SaSSy proteins are both typically located in the cytoplasm of WT plants (Fig. 1A and B), whereas after fusion with LTP, the GFP signal of LTP-SaSSy was observed in chloroplasts. In contrast, the LTP-FPS protein appeared in both cytoplasm and chloroplasts (Fig. 1C and D). Compared with the localization of LTP-SaSSy and LTP-FPS proteins, the GFP signals of RTP-FPS and RTP-SaSSy were exclusively visible in the chloroplasts (Fig. 2E and F). These observations indicated that the targeting efficiency of RTP is higher than that of the LTP, as it completely conveyed both FPS and SaSSy proteins into the chloroplasts. To further study the effects of co-expression of RTP-FPS and RTP-SaSSy on the production of santalenes and bergamotene, these genes were over-expressed in both *N. benthamiana* and *N. tabacum*.

**Figure 1.**
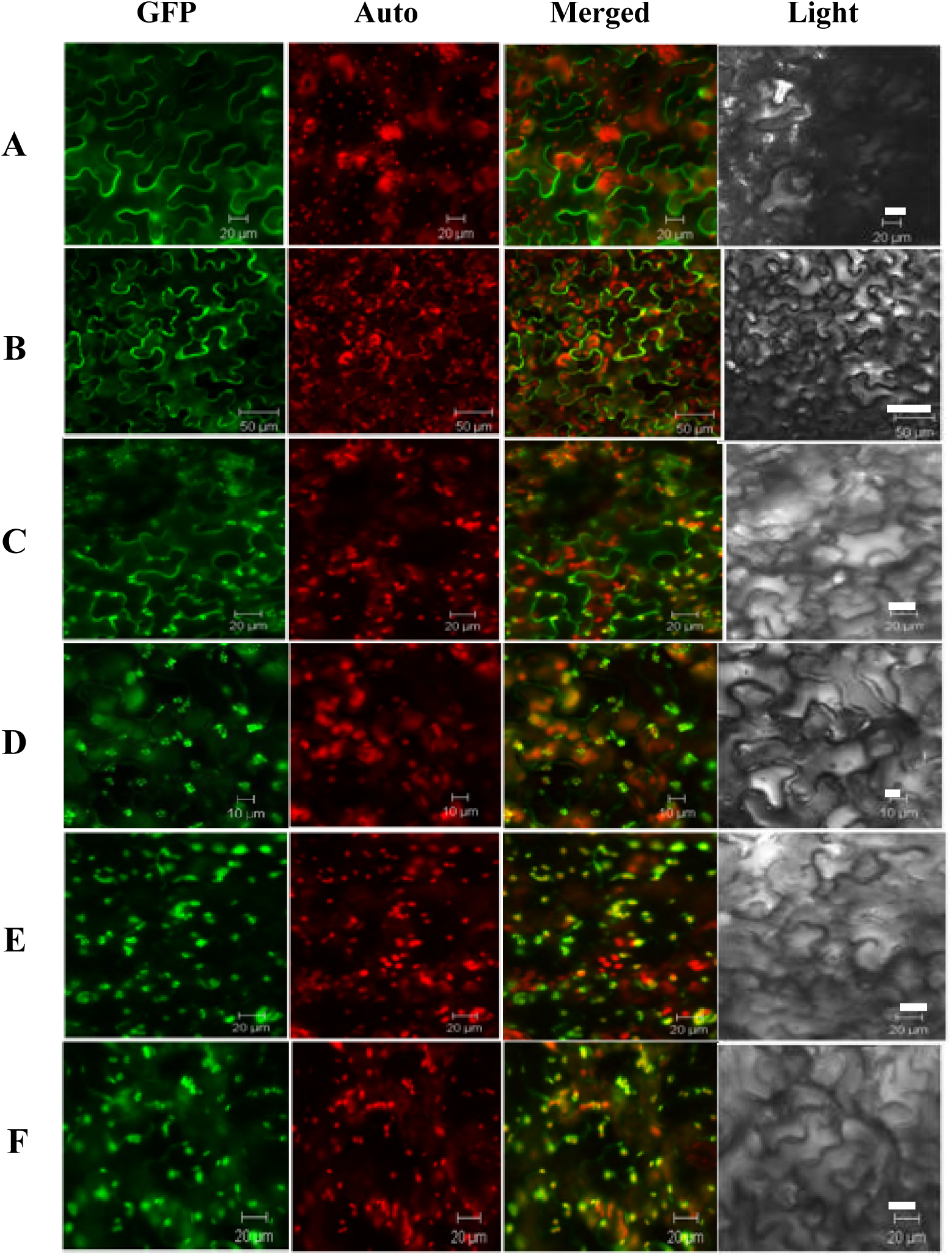
Localization of TP-FPS-GFP, TP-SaSSy-GFP, FPS-GFP, and SaSSy-GFP proteins in the leaves of *Nicotiana benthamiana*. **(A)** FPS-GFP located in the cytoplasm, scale bar = 20 µm; **(B)** SaSSy-GFP located in the cytoplasm, scale bar = 50 µm; **(C)** LTP-FPS-GFP located in both cytoplasm and chloroplasts, scale bar = 20 µm; **(D)** LTP-SaSSy-GFP located in the chloroplasts, scale bar = 10 µm; **(E)** RTP-FPS-GFP located in the chloroplasts, scale bar = 20 µm; **(F)** RTP-SaSSy-GFP located in the chloroplasts, scale bar = 20 µm. LTP, *Solanum lycopersicum* linalool synthase transit peptide; RTP, *Arabidopsis thaliana* ribulose bisphosphate carboxylase small subunit transit peptide; SaSSy, *Santalum album* santalene synthase; FPS, *A*. *thaliana* farnesyl diphosphate synthase; GFP, green fluorescent protein.

**Figure 2.**
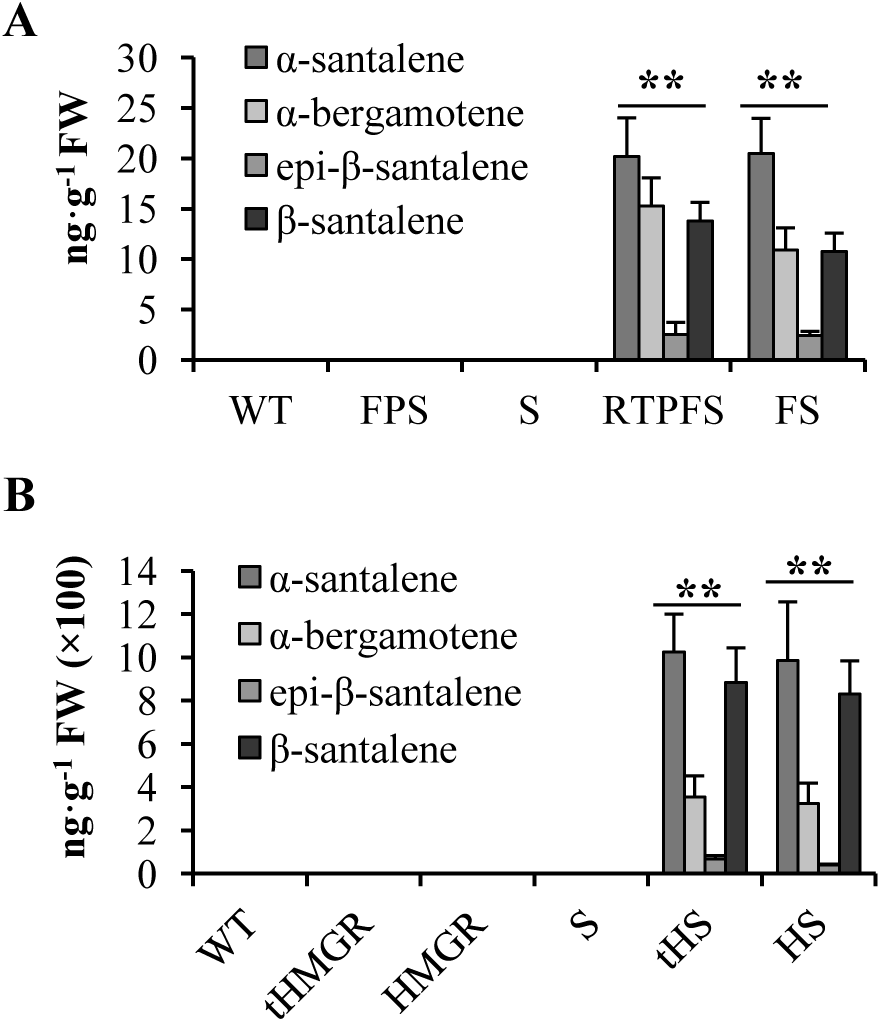
Chloroplast targeting and MVA enhancement strategies enhance the production of santalenes and bergamotene in *Nicotiana benthamiana* agro-infiltrated leaves. **(A)** The production of santalenes and bergamotene in *N. benthamiana* transiently expressing *RTP*-*FPS* + *RTP*-*SaSSy* or *FPS* + *SaSSy*; **(B)** The production of santalenes and bergamotene in *N. benthamiana* transiently expressing *tHMGR* + *SaSSy* or *HMGR* + *SaSSy*. RTP, *Arabidopsis thaliana* ribulose bisphosphate carboxylase small subunit transit peptide; SaSSy, *Santalum album* santalene synthase; FPS, *A*. *thaliana* farnesyl diphosphate synthase; HMGR, *A*. *thaliana* 3-hydroxy-3-methylglutaryl-CoA reductase; tHMGR, *A*. *thaliana* truncated HMGR; WT, wild type; S, lines expressing *SaSSy*; FS, lines transiently expressing *FPS* + *SaSSy*; RTPFS, lines transiently expressing *RTP*-*FPS* + *RTP*-*SaSSy*; tHS, lines transiently expressing *tHMGR* + *SaSSy*; HS, lines transiently expressing *HMGR* + *SaSSy*. The mean ± SD values of three biological replicates are shown. Asterisks indicate statistically significant differences of total santalenes and bergamotene from S based on Student’s unequal variance *t*′-test (* P < 0.05 and ** P < 0.01).

### Effects of different metabolic engineering methods on the production of santalenes and bergamotene in *N. benthamiana*

To assess the effects of chloroplast targeting and MVA enhancement strategies using *A. thaliana* FPS and HMGR or tHMGR on the production of santalenes and bergamotene, we transiently co-expressed the two genes together in *N. benthamiana* (*RTP*-*FPS* + *RTP*-*SaSSy, FPS* + *SaSSy, HMGR* + *SaSSy,* or *tHMGR* + *SaSSy*). To avoid gene silencing mediated by promoter homology [33], the *SaSSy* or *RTP*-*SaSSy* genes were placed under the control of the *RbcS* promoter of *A. thaliana*, whereas each of the *RTP*-*FPS, FPS, tHMGR,* and *HMGR* sequences was expressed under the control of the Cauliflower mosaic virus (CaMV) 35S promoter.

Three days after *N. benthamiana* agro-infiltration, 200 mg of treated leaves from each plant was ground in liquid nitrogen and extracted using ethyl acetate, and the resulting extracts were analyzed by GC–MS. Although no santalenes or bergamotene were observed in the wild-type *N. benthamiana* plant and *N. benthamiana* leaves transiently expressing a single gene (*SaSSy*) (Fig. 2A), trace amounts of santalenes and bergamotene were found in lines transiently co-expressing *RTP*-*FPS* + *RTP*-*SaSSy* or *FPS* + *SaSSy* (Fig. 2A), which indicated that the chloroplast targeting and FPS enhancement strategies slightly increased the production of santalenes and bergamotene. When *tHMGR* was transiently co-expressed with *SaSSy*, the tobacco plants accumulated 1025.8 ng·g^-1^ fresh weight (FW) of *α*-santalene, 354.8 ng·g^-1^ FW of *α*-bergamotene, 68.8 ng·g^-1^ FW of *Epi-β-*santalene, and 885.1 ng·g^-1^ FW of *β*-santalene, the values of which were approximately 50 times higher than the amounts produced using the chloroplast targeting and FPS enhancement strategies (Fig. 2B). In addition, co-expression with *HMGR* also yielded high amounts of santalenes and bergamotene: 984.8 ng·g^-1^ FW of *α*-santalene, 324.1 ng·g^-1^ FW of *α*-bergamotene, 40.5 ng·g^-1^ FW of *Epi-β*-santalene, and 830.0 ng·g^-1^ FW of *β*-santalene (Fig. 2B). Therefore, the *HMGR* and *tHMGR* MVA enhancement strategy is suggested to be a more suitable method for enhancing the yields of santalenes and bergamotene in *N. benthamiana*. To further evaluate the effects of the different metabolic engineering methods on the production of santalenes and bergamotene, we transformed *N. tabacum* plants with *RTP*-*FPS* + *RTP*-*SaSSy, FPS* + *SaSSy, tHMGR* + *SaSSy,* or *HMGR* + *SaSSy*.

### HMGR and tHMGR enhance the production of santalenes and bergamotene in transgenic *N. tabacum*

The GC–MS results of *N. benthamiana* agro-infiltration experiments indicated that chloroplast targeting and co-expression of *FPS* and *HMGR* or *tHMGR* improved the production of santalenes and bergamotene to different levels. To further investigate the effects of these metabolic engineering methods on the production of santalenes and bergamotene in stable transgenic *N. tabacum* plants, different gene combinations (*SaSSy, RTP*-*FPS* + *RTP*-*SaSSy, FPS* + *SaSSy, tHMGR* + *SaSSy*, or *HMGR* + *SaSSy*) were used to transform *N. tabacum*. For each transformation event, three independent lines with similar relative expression levels of *SaSSy* were selected from at least 10 lines of transgenic plants and GC–MS was performed to analyze the volatiles produced by each of the selected transgenic plants.

No santalenes or bergamotene were observed in the wild-type *N. tabacum* plant (Fig. 3B and D). Three transgenic lines (S3, S4, and S16), which had comparable relative transcript level of *SaSSy* (Fig. S2 and Fig. 3A), produced 162.6–241.8 ng·g^-1^ FW·24 h^-1^ of *α*-santalene, 17.1–25.5 ng·g^-1^ FW·24 h^-1^ of *α*-bergamotene, 11.5–18.3 ng·g^-1^ FW·24 h^-1^ of *epi*-*β*-santalene, and 73.9–166.6 ng·g^-1^ FW·24 h^-1^ of *β*-santalene (Table 2 and Fig.3B). Additionally, three transgenic lines (RTPFS8, 10, and 15) expressing *RTP*-*FPS* + *RTP*-*SaSSy* (S2 Fig. and Fig. 3A), each of which had relative expression levels of *SaSSy* similar to those of S3, 4, and 16, produced 85.1–112.2 ng·g^-1^ FW·24 h^-1^ of *α*-santalene, 11.6–19.1 ng g^-1^ FW·24 h^-1^ of *α*-bergamotene, 14.1–45.7 ng g^-1^ FW·24 h^-1^ of *epi-β*-santalene, and 58.7–82.2 ng·g^-1^ FW·24 h^-1^ of *β*-santalene (Table 2 and Fig. 3B). These results showed that the average production of santalenes and bergamotene in RTPFS8, 10, and 15 was slightly lower than that of S3, S4, and S16.

**Table 1.** Yields of santalenes and bergamotene in *Nicotiana benthamiana* using agro-infiltration (ng·g^-1^ fresh weight)

| | α-santalene | α-bergamotene | epi-β-santalene | β-santalene |
| --- | --- | --- | --- | --- |
| WT<sup>a</sup> | nd<sup>b</sup> | nd | nd | nd |
| FPS | nd | nd | nd | nd |
| tHMGR | nd | nd | nd | nd |
| HMGR | nd | nd | nd | nd |
| S<sup>c</sup> | nd | nd | nd | nd |
| RTPFS<sup>d</sup> | 20.2 ± 10.8<sup>e</sup> | 15.2 ± 4.7 | 2.5 ± 1.1 | 13.8 ± 6.8 |
| FS<sup>f</sup> | 20.4 ± 5.5 | 10.9 ± 4.2 | 2.4 ± 0.4 | 10.7 ± 3.8 |
| tHS<sup>g</sup> | 1025.8 ± 374.1 | 354.8 ± 97.6 | 68.8 ± 15.0 | 885.1 ± 258.7 |
| HS<sup>h</sup> | 984.8 ± 572.4 | 324.1 ± 94.9 | 40.5 ± 4.4 | 830.0 ± 254.2 |

<sup>a</sup> WT, wild type; <sup>b</sup> nd, not detected; <sup>c</sup> S, lines expressing *SaSSy*; <sup>d</sup> RTPFS, lines expressing *RTPFPS* + *RTPSaSSy*; <sup>e</sup> ± , mean ± SD, n = 3; <sup>f</sup> FS, lines expressing *FPS* + *SaSSy*; <sup>g</sup> tHS, lines expressing *tHMGR* + *SaSSy*; <sup>h</sup> HS, lines expressing *HMGR* + *SaSSy*

**Table 2.** Yields of santalenes and bergamotene in transgenic *Nicotiana tabacum* (ng·g^-1^·fresh weight 24 h^-1^)

| | α-santalene | α-bergamotene | epi-β-santalene | β-santalene |
| --- | --- | --- | --- | --- |
| WT<sup>a</sup> | nd<sup>b</sup> | nd | nd | nd |
| S3<sup>c</sup> | 172.2 ± 95.2<sup>d</sup> | 17.1 ± 6.3 | 18.3 ± 9.7 | 73.9 ± 16.2 |
| S4 | 241.8 ± 97.8 | 25.5 ± 12.5 | 17.3 ± 10.2 | 166.6 ± 75.2 |
| S16 | 162.6 ± 78.9 | 18.7 ± 9.8 | 11.5 ± 5.5 | 77.7 ± 36.6 |
| RTPFS8<sup>e</sup> | 112.2 ± 66.3 | 11.6 ± 6.6 | 15.5 ± 5.6 | 58.7 ± 32.2 |
| RTPFS10 | 85.1 ± 62.5 | 15.7 ± 9.8 | 14.1 ± 7.7 | 71.7 ± 32.3 |
| RTPFS15 | 91.4 ± 23.5 | 19.1 ± 9.8 | 45.7 ± 31.1 | 82.2 ± 25.4 |
| FS1<sup>f</sup> | 114.4 ± 62.5 | 54.1 ± 22.1 | 23.2 ± 9.8 | 98.4 ± 36.7 |
| FS10 | 98.7 ± 26.5 | 19.1 ± 10.2 | 6.4 ± 1.5 | 21.9 ± 9.7 |
| FS15 | 315.3 ± 99.9 | 26.4 ± 12.1 | 7.3 ± 3.4 | 169.7 ± 74.2 |
| tHS10<sup>g</sup> | 557.2 ± 123.3 | 21.3 ± 10.2 | 3.1 ± 0.9 | 314.3 ± 111.6 |
| tHS11 | 789.1 ± 321.2 | 45.5 ± 31.2 | 142.2 ± 46.5 | 951.3 ± 351.2 |
| tHS13 | 1125.1 ± 366.6 | 114.3 ± 35.6 | 207.7 ± 99.8 | 1275.7 ± 625.5 |
| HS4<sup>h</sup> | 537.1 ± 177.5 | 62.3 ± 24.1 | 53.3 ± 39.8 | 207.7 ± 99.9 |
| HS5 | 495.7 ± 111.9 | 52.2 ± 18.9 | 31.7 ± 12.1 | 312.1 ± 88.8 |
| HS6 | 2946.1 ± 693.8 | 268.6 ± 87.8 | 294.2 ± 98.8 | 1732.3 ± 732.9 |

<sup>a</sup> WT, wild type; <sup>b</sup> nd, not detected; <sup>c</sup> S, lines expressing *SaSSy*; <sup>d</sup> RTPFS, lines expressing *RTPFPS* + *RTPSaSSy*; <sup>e</sup> ± , mean ± SD, n = 3; <sup>f</sup> FS, lines expressing *FPS* + *SaSSy*; <sup>g</sup> tHS, lines expressing *tHMGR* + *SaSSy*; <sup>h</sup> HS, lines expressing *HMGR* + *SaSSy*

**Figure 3.**
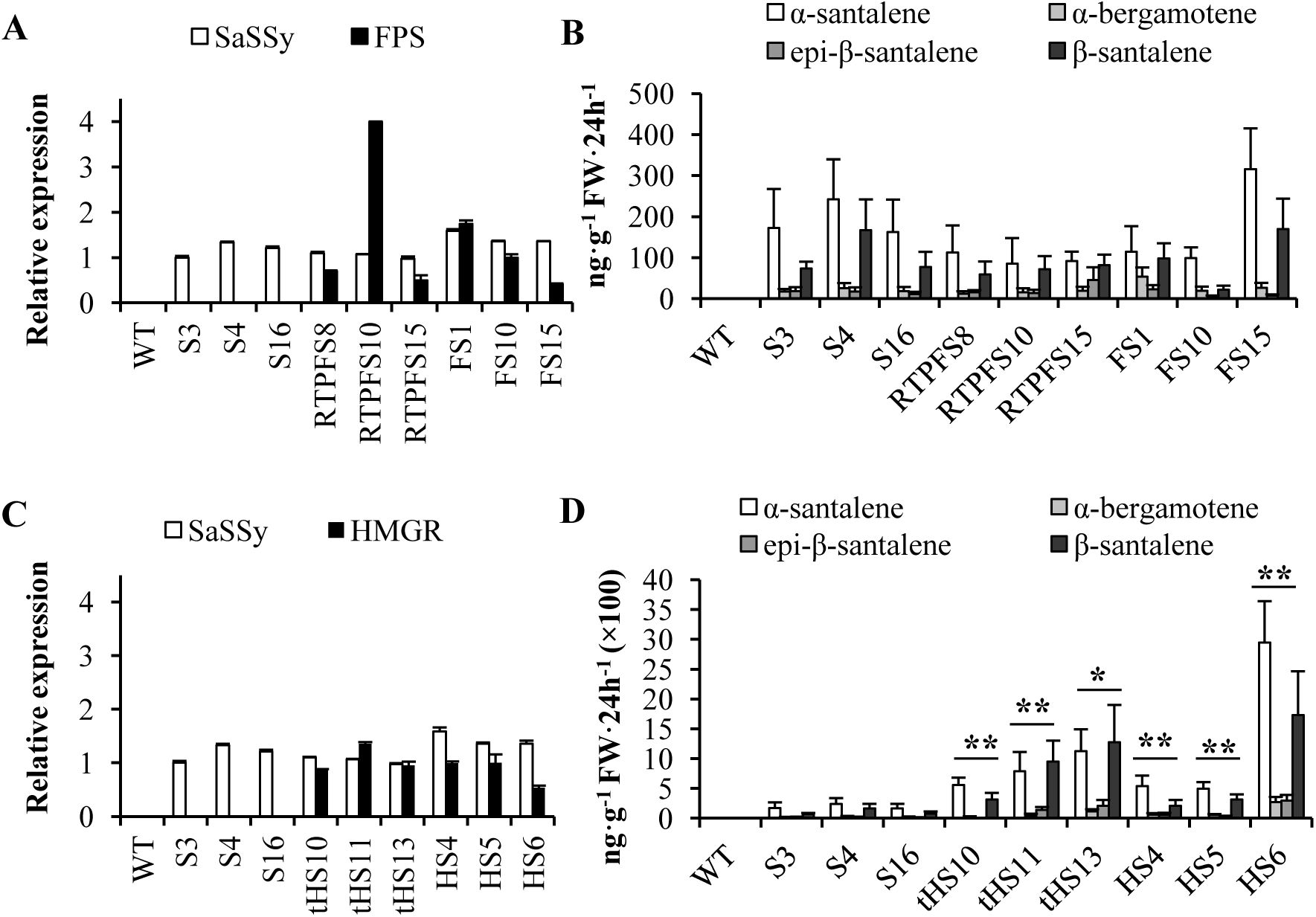
The HMGR and tHMGR MVA enhancement strategy enhances the production of santalenes and bergamotene in transgenic *Nicotiana tabacum.* **(A)** Relative transcript level analysis of transgenic *N. tabacum* expressing *RTP*-*FPS* + *RTP*-*SaSSy* or *FPS* + *SaSSy*; **(B)** Production of santalenes and bergamotene in transgenic *N. tabacum* expressing *RTP*-*FPS* + *RTP*-*SaSSy* or *FPS* + *SaSSy*; **(C)** Relative transcript level analysis of transgenic *N. tabacum* expressing *tHMGR* + *SaSSy* or *HMGR* + *SaSSy*; **(D)** The production of santalenes and bergamotene in transgenic *N. tabacum* expressing *tHMGR* + *SaSSy* or *HMGR* + *SaSSy*. RTP, *Arabidopsis thaliana* ribulose bisphosphate carboxylase small subunit transit peptide; SaSSy, *Santalum album* santalene synthase; FPS, *A*. *thaliana* farnesyl diphosphate synthase; HMGR, *A*. *thaliana* 3-hydroxy-3-methylglutaryl-CoA reductase; tHMGR, *A*. *thaliana* truncated HMGR. *EF1A-1* was used as a control gene. WT, wild type; S, lines expressing *SaSSy*; FS, lines expressing *FPS* + *SaSSy*; RTPFS, lines expressing *RTP*-*FPS* + *RTP*-*SaSSy*; tHS, lines expressing *tHMGR* + *SaSSy*; HS, lines expressing *HMGR* + *SaSSy*. The mean ± SD values of three biological replicates are shown. Asterisks indicate statistically significant differences of total santalenes and bergamotene from S4 based on Student’s unequal variance *t*′-test (* P < 0.05 and ** P < 0.01).

Using the same method, FS1, 10, and 15; tHS10, 11, and 13; and HS4, 5, and 6 lines were selected from corresponding transgenic plants (S2 Fig. and Fig. 3C). The FS1, 10, and 15 lines produced approximately 98.7–315.3 ng·g^-1^ FW·24 h^-1^ of *α*-santalene, 19.1–54.1 ng·g^-1^ FW·24 h^-1^ of *α*-bergamotene, 6.4–23.2 ng·g^-1^ FW·24 h^-1^ of *epi-β*-santalene, and 21.9–169.7 ng·g^-1^ FW·24 h^-1^ of *β*-santalene (Table 2 and Fig. 3B). The average production of santalenes and bergamotene in FS1, 10 and 15 was similar to that of S3, S4, and S16 plants. Furthermore, tHS10, 11, and 13 lines produced 557.2–1125.1 ng·g^-1^ FW·24 h^-1^ of *α*-santalene, 21.3–114.3 ng·g^-1^ FW·24 h^-1^ of *α*-bergamotene, 3.1–207.7 ng g^-1^ FW·24 h^-1^ of *epi-β*-santalene, and 314.3–1275.7 ng g^-1^ FW·24 h^-1^ of *β*-santalene, whereas the HS4, 5, and 6 lines produced 495.7–2946.1 ng g^-1^ FW·24 h^-1^ of *α*-santalene, 52.2–268.6 ng·g^-1^ FW·24 h^-1^ of *α*-bergamotene, 31.7–294.2 ng g^-1^ FW·24 h^-1^ of *epi-β*-santalene, and 207.7–1732.3 ng g^-1^ FW·24 h^-1^ of *β*-santalene (Table 2 and Fig. 3D). These results revealed that co-expression of *FPS* with either *HMGR* or *tHMGR* produced 500 to 1000 times more santalenes and bergamotene than the S3, S4, and S16 lines, indicating that *HMGR* or *tHMGR* could substantially enhance the carbon flux within the MVA pathway in *N. tabacum* plants, which was thereby available for santalene and bergamotene biosynthesis. The GC–MS chromatography and MS data for transgenic tobacco headspace samples are shown in S3 Fig.

### Production of santalols and bergamotol in transgenic tobacco

Santalols and bergamotol, which are major components of sandalwood essential oil, are generated from santalenes and bergamotene by *S. album* santalene/bergamotene CYP76F cytochrome P450 [23]. The results of the present study indicated that co-expression of HMGR in transgenic tobacco could enhance the production of santalenes and bergamotene. We subsequently employed *HMGR, SaSSy,* and *S. album* santalene/bergamotene oxidase *SaCYP76F39v1* [23] to produce santalols and bergamotol in tobacco. To avoid gene silencing caused by promoter homology [33], three strong constitutive promoters, Arabidopsis *ubiquitin 10* [27], *RbcS*, and 35S promoters were used to control these three genes respectively. Unfortunately, only santalenes and bergamotene were detected in both agro-infiltrated *N. benthamiana* plants and transgenic *N. tabacum* plants co-expressing *HMGR, SaSSy,* and *SaCYP76F39v1*) (S3 Fig.), the relative gene expression levels of which were analyzed by qRT-PCR (S2 Fig.).

To detect the presence of glycosylated santalols and bergamotol in transgenic tobacco plants, homogenized HSP1 leaves were treated with either a saturated CaCl_2_ solution or a glycosidase [34]. In the present study, we were unable to detect any santalols or bergamotol in either untreated or treated leaves of HSP1 line plants (S4 Fig.). These results indicate that santalols and bergamotol were not stored as nonvolatile santalols/bergamotol–glycoside forms in leaf tissues or cells of transgenic tobacco plants.

### Interaction of aphids with transgenic *N. tabacum* emitting santalenes and bergamotene

Green peach aphids are major pests of many commercial crops, not only causing feeding damage but also transmitting various viruses that can result in substantial losses [35].

To investigate the interaction of aphids with transgenic *N. tabacum* plants emitting santalenes and bergamotene, two-choice experiments were performed to examine green peach aphid preference between WT and transgenic tobacco plants [36]. Initially, we conducted short-term two-choice experiments using plants placed in a cage (Fig. 4A), which revealed that 59.8% to 63.6% of aphids showed a preference for the transgenic tobacco plants (S3, 4, and 16), indicating that the transgenic plants emitting santalenes and bergamotene tended to attract aphids to a greater extent than did the WT plants. Subsequently, to examine the long-term preference of insects for either WT or transgenic tobacco plants. Four weeks after releasing aphdis, we observed that 67.1% to 67.3% of aphids showed preference for transgenic tobacco (S3, 4, and 16; Fig. 4B and D, S5 Fig.). These results indicated that the transgenic plants emitting santalenes and bergamotene were more attractive to aphids than were WT plants over the long-term. To further confirm the aphid-attractant properties of santalenes and bergamotene, solutions of these terpenes were also utilized in two-choice tests using agar medium plates. Consistent with the transgenic tobacco choice tests, the leaves of wild-type *N. tabacum* plants treated with santalene and bergamotene solutions also proved to be more attractive to green peach aphids (Fig. 4C and S6 Fig.).

**Figure 4.**
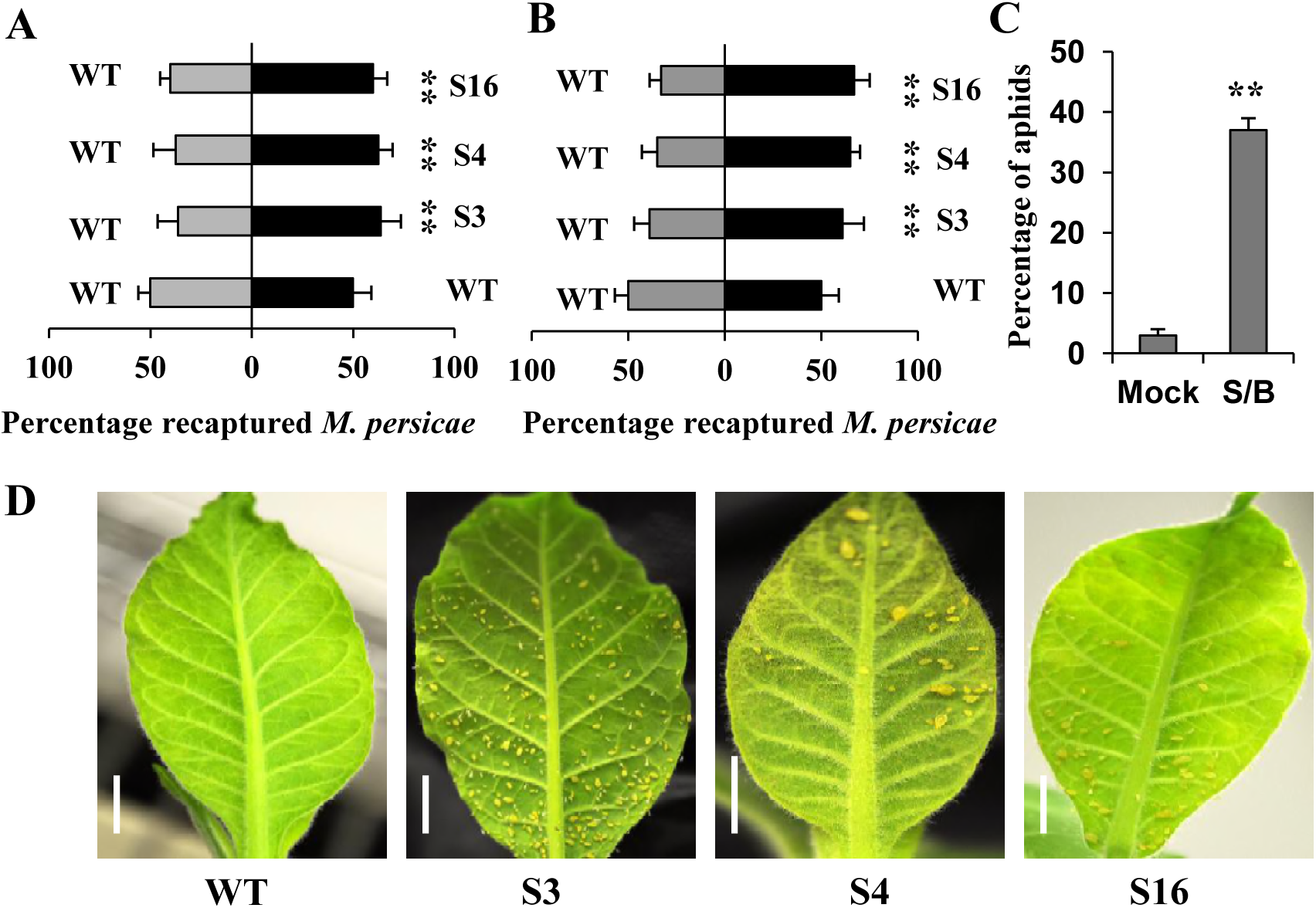
Green peach aphid choice experiments. **(A)** Transgenic tobacco lines emitting santalenes and bergamotene attracted aphids over the short-term in two-choice experiments (n = 8); **(B)** Long-term choice experiments for aphids using the transgenic lines and WT tobacco in a growth chamber (n = 8); **(C)** Green peach aphid two-choice experiments using agar medium plates treated with santalene and bergamotene solutions and hexane (n = 8); **(D)** Aphids on the leaves of transgenic line and WT tobacco in the long-term choice experiment, scale bar = 1 cm. WT, wild type; S, lines expressing *SaSSy*; Mock, wild-type *Nicotiana tabacum* leaves treated with hexane; S/B, wild-type *N. tabacum* leaves treated with santalene and bergamotene solution. The mean ± SD values for eight biological replicates are shown. Asterisks indicate statistically significant differences from the WT based on Student’s *t*-test (* P < 0.05 and ** P < 0.01).

### Discussion

*N. benthamiana* transient agro-infiltration and generation of transgenic *N. tabacum* plants were used to assess the effects of different methods (chloroplast targeting and co-expression of *FPS* and *HMGR* or *tHMGR*) on the production of santalenes and bergamotene and santalols and bergamotol; thereafter, we investigated the interaction of aphids with santalenes and bergamotene based on aphid two-choice experiments. We demonstrated that the HMGR or tHMGR MVA pathway enhancement strategy was the simplest and most effective approach for generating santalenes and bergamotene in tobacco; however, the santalenes and bergamotene thus produced could not be further converted to santalols and bergamotol by santalene/bergamotene oxidase. In addition, using choice experiments, we demonstrated that aphids were strongly attracted to transgenic tobacco plants emitting santalenes and bergamotene, indicating that such plants have potential application as a trap crop for pest management.

### Biosynthesis of santalenes and bergamotene using different metabolic engineering methods

The results of *N. benthamiana* agro-infiltration and transgenic *N. tabacum* generation using the chloroplast targeting method indicated that this method is not effective in enhancing the synthesis of santalenes and bergamotene. Evidence tended to indicate that chloroplast targeting for santalene and bergamotene biosynthesis is too complex and inefficient.

LTPFPS protein was observed in both cytosol and chloroplasts, indicating that the LTP could not completely convey the FPS protein into chloroplasts (Fig. 1C). This result indicates that prior to generating transgenic tobacco plants, additional confocal laser screening is necessary to confirm the targeting efficiency of different transit peptides, as targeting efficiency tends to be affected by the structure and sequence motifs of the transit peptides and sesquiterpene synthases [32].

Monoterpenes or sesquiterpenes were not detected in the volatiles of WT plants, which could be attributed to the very low contents of IPP and DMAPP available for the generation of secondary metabolite volatile terpenes in normal WT tobacco leaves (S3 Fig.). When RTPFS and RTPSaSSy were over-expressed in *N. benthamiana* (Fig. 2A) and transgenic *N. tabacum* (RTPFS8, RTPFS10, and RTPFS15; Fig. 3B), trace amounts of both santalenes and bergamotene were detected, which may be due to the low levels of chloroplastic IPP and DMAPP. Since 1-deoxy-D-xylulose-5-phosphate synthase (DXS) has been suggested to mediate a rate-limiting step in the chloroplastic MEP pathway [37], we hypothesize that if *DXS* was co-expressed with *RTPFPS* and *RTPSaSSy*, a greater carbon flux for santalene and bergamotene biosynthesis could have been generated via the MEP pathway.

HMGR is known to catalyze an important rate-limiting step in the MVA pathway of plants [16]. Furthermore, tHMGR, which is believed to be a soluble non-feedback-regulated rate-limiting enzyme in this pathway, has a better enhancement effect [38]. Therefore, Arabidopsis HMGR and tHMGR were utilized in anticipation of enhancing activity of the MVA pathway in tobacco. We observed, however, that both HMGR and tHMGR had similar enhancement effects on the synthesis of santalenes and bergamotene (Fig. 2B and 3D); thus, tHMGR did not appear to confer any advantage in this respect compared with HMGR. HMGR activity is regulated not only by endoplasmic reticulum (ER)-associated degradation [39] but also is controlled by AMP-dependent protein kinase-catalyzed phosphorylation [40]. Although tHMGR could conceivably prevent the regulation of ER-associated degradation due to truncation of the ER signal peptide, it might be regulated by AMP kinase similarly to HMGR. We hypothesize that when energy is consumed by the activity of the MVA pathway, which was enhanced by over-expression of tHMGR to produce a higher amount of santalenes and bergamotene, the levels of AMP kinase in energy-depleted cells increase and subsequently inactivate tHMGR. As a consequence, the transgenic tobacco plants expressing tHMGR produce amounts of santalenes and bergamotene that are similar to those produced by the HMGR-expressing lines.

Therefore, before transformation of plant metabolic engineering, it is necessary to use GC-MS to analyze the volatile components of plants and to elucidate the expression of the MVA and MEP pathways in plants, which can effectively guide us to adopt appropriate metabolic engineering programs.

Although *SaCYP76F39v1* cloned from *S. album* santalene/bergamotene oxidases [23] was jointly co-expressed with *SaSSy* and *HMGR* in tobacco, we were unable to detect santalols and bergamotol in the agro-infiltrated *N. benthamiana* and transgenic *N. tabacum*, indicating that *SaCYP76F39v1* could not hydroxylate santalenes and bergamotene to form santalols and bergamotol in tobacco. Capitate glandular trichomes are the main site of aromatic compound synthesis in tobacco; however, unlike peltate glandular trichomes, the absence of stratum corneum spaces dictates that although capitate glandular trichomes can synthesize terpenes, they are unable to store these compounds [41]. Hence, we hypothesize that any santalene and bergamotene synthesized might volatilize from leaves before they can be converted to santalols and bergamotol by *SaCYP76F39v1*. Alternatively, it is believed that *SaCYP76F39v1* may have low catalytic activity in tobacco cells.

### Attraction of aphids to santalenes/bergamotene

Insects can locate their host plants by detecting specific olfactory signals. Plant volatiles play an important role in this host-location process, and herbivorous insects can distinguish certain host plants from the ratio-specific blends of specific volatiles produced by these plants [42].

Several types of learning mechanisms, such as habituation learning, associative learning, aversion learning, sensitization, and induction of preference, have been identified in phytophagous insects [43]. In the short-term two-choice experiment conducted in the present study, we found that when given the choice between transgenic and WT tobacco plants, green peach aphids show a preference for the former. Similar results were obtained in long-term choice experiments, with most green peach aphids showing a preference for transgenic tobacco (Fig. 4B and C), thereby indicating the operation of some form of recognition and learning mechanism in these aphids. We inferred that green peach aphids were attracted to transgenic tobacco at the beginning of the experiment by the santalene/bergamotene odors emitted by these plants, and that after several generations, the aphid progeny had adapted to transgenic tobacco, such that they might associate santalene/bergamotene odors with host plants via associative learning. Hence, in the long-term choice experiments, when aphids migrated to new leaves, they chose the santalene- and bergamotene-emitting transgenic tobacco rather than WT tobacco.

*trans*-*α*-Bergamotene has been identified as an active compound in the sex attractant recognized by male *Melittobia digitata* [44], and may play a similar important role in the interaction with green peach aphids. However, given that purified standards of *α*-santalene, *trans*-*α*-bergamotene, epi-*β*-santalene, and *β*-santalene are currently commercially unavailable, this somewhat limits our scope for further investigating the interaction of each of these compounds with aphids.

The use of trap crops is one of the most effective pesticide-free pest management strategies, in which phytophagous pests are typically attracted away from the main crop [45]. Green peach aphids are very common and a source of considerable damage to agricultural plants. To combat these pests, employment of trap crops is one of the several major control methods [46]. Our short- and long-term choice experiments demonstrate that transgenic tobacco can potentially be deployed to serve as a trap crop for green peach aphid management.

In conclusion, we utilized an optimized HMGR or tHMGR MVA pathway strategy to generate higher amounts of santalenes and bergamotene in transgenic tobacco plants. However, the santalenes and bergamotene thus produced could not be further hydroxylated to santalols and bergamotol by *S. album* santalene/bergamotene oxidase. In both short- and long-term choice experiments, green peach aphids were strongly attracted to transgenic tobacco lines emitting santalenes/bergamotene, indicating that such lines could be employed as a trap crop to prevent green peach aphid infestation in wild-type tobacco. In the future, we plan to conduct field trials to assess the potential utility of these transgenic lines in pest management.

## Acknowledgements

This work was financially supported by a grant from the Singapore National Research Foundation (Competitive Research Programme Award No. NRF-CRP8-2011-02), Natural Science Foundation of China (No. 21768005) and the Temasek Life Sciences Laboratory. We wish to thank Professor Ji Lianghui for provision of santalene and bergamotene solutions.

## Author contributions statement

J.L.Y. conceived the experiments, analyzed the results, and wrote the main manuscript text. J.L.Y. and W.S.W. conducted the experiments. Both authors reviewed the manuscript.

## Conflict of interest

The authors declare that they have no competing interests.

## Supporting information

Additional Supporting information data associated with this article can be found in the online version:

**S1 Table. Oligonucleotides used in vector construction.**

**S2 Table. Gene names and GenBank reference numbers.**

**S3 Table. Primers used in qPCR.**

**S1 Figure. Vector construction for santalol biosynthesis.**

**S2 Figure. Relative transcript level analysis of transgenic *Nicotiana tabacum*.**

**S3 Figure. GC–MS chromatograms of transgenic *Nicotiana tabacum* headspace volatiles.**

**S4 Figure. GC–MS analysis of glycosylated santalols/bergamotol in transgenic tobacco (HSP1).**

**S5 Figure. Green peach aphid long-term choice experiments using santalene- and bergamotene-emitting transgenic *Nicotiana tabacum* plants.**

**S6 Figure. Green peach aphids two-choice experiments on agar medium plates using santalene and bergamotene solutions: scale bar = 1 cm.**

## Abbreviations

ADS: amorpha-4,11-diene synthase
DMAPP: dimethyla llyl diphosphate
FPS: farnesyl diphosphate synthase
FPP: farnesyl diphosphate
HMGR: 3-hydroxy-3-methylglutaryl-CoA reductase
IPP: isopentenyl diphosphate
GC–MS: gas chromatography–mass spectrometry
*LTP*: linalool synthase transit peptide
MVA: mevalonic-acid
*RTP,*: RbcS secretion signal
SaSSy: santalene synthase
tHMGR: truncated 3-hydroxy-3-methylglutaryl-CoA reductase
WT: wild-type

